# Reactive SINDy: Discovering governing reactions from concentration data

**DOI:** 10.1101/442095

**Authors:** Moritz Hoffmann, Christoph Fröhner, Frank Noé

## Abstract

The inner workings of a biological cell or a chemical reaction can be rationalized by the network of reactions, whose structure reveals the most important functional mechanisms. For complex systems, these reaction networks are not known a priori and cannot be efficiently computed with *ab initio* methods, therefore an important approach goal is to estimate effective reaction networks from observations, such as time series of the main species. Reaction networks estimated with standard machine learning techniques such as least-squares regression may fit the observations, but will typically contain spurious reactions. Here we extend the sparse identification of nonlinear dynamics (SINDy) method to vector-valued ansatz functions, each describing a particular reaction process. The resulting sparse tensor regression method “reactive SINDy” is able to estimate a parsimonious reaction network. We illustrate that a gene regulation network can be correctly estimated from observed time series.

## I. INTRODUCTION

Mapping out the reaction networks behind biological processes, such as gene regulation in cancer [1], is paramount to understanding the mechanisms of life and disease. A well-known example of gene regulation is the lactose operon whose crystal structure was resolved in [24] and dynamics were modeled in [43]. The system’s “combinatorial control” in E. coli cells was quantitatively investigated in [22], in particular studying repression and activation effects. These gene regulatory effects often appear in complex networks [35] and there exist databases resolving these for certain types of cells, e.g., E. coli cells [11] and yeast cells [23]. Another example where mapping the active reactions is important is that of chemical reactors [30], where understanding which reactions are accessible for a given set of educts and reaction conditions is important to design synthesis pathways [7, 20].

The traditional approach to determine a reaction network is to propose the structure of the network based on chemical insight and subsequently fit the parameters given available data [32]. To decipher complex reaction environments such as biological cells, it would be desirable to have a data-driven approach that can answer the question which reactions are underlying a given observation, e.g., the time series of a set of reactants. However, in sufficiently complex reaction environments the number of reactive species and possible reactions is practically unlimited – as an illustration, consider vast amount of possible isomerizations and post-translational modifications for a single protein molecule. Therefore, the more specific formulation is “given observations of a set of chemical species, what is the *minimal set* of reactions necessary to explain their time evolution?”. This formulation calls for a machine learning method that can infer the reaction network underlying the observation data.

Knowledge about the reaction network can be applied to parameterize other numerical methods to further investigate the processes at hand. Such methods include particle-based approaches derived from the chemical master equation [13, 18, 41, 42], as well as highly detailed but parameter-rich methods such as particle-based or interacting-particle reaction dynamics [2, 8, 10, 17, 33, 39, 40] capable of fully resolving molecule positions in space and time – see [3, 34] for recent reviews.

Existing methods to infer regulatory networks include ARCANE [26] that uses experimental essay data and information theory, as well as the likelihood approach presented in [37] that takes the stochasticity of observed reactant time series into account.

The method presented in this work can identify underlying complex reaction networks from concentration time series by following the law of parsimony, i.e., by inducing sparsity in the resulting reaction network. This promotes the interpretability of the model and avoids overfitting. We formulate the problem as data-driven identification of a dynamical system, which renders the method consistent with and an extension of the framework of sparse identification of nonlinear dynamics (SINDy) [5]. Specifically, the problem of identifying a reaction network from time traces of reactant concentrations can be solved by finding a linear combination from a library of nonlinear ansatz functions that each corresponds to a reaction acting on a set of reactants. With this formulation, the reaction rates can be determined *via* regression. Sparsity is induced by equipping the regression algorithms with a sparsity inducing regularization. SINDy was investigated, generalized, and applied in many different ways, e.g., including control [6] (SINDYc), in the context of partial differential equations [31], updating already existing models [29] (abrupt-SINDy), and looking into convergence properties [44].

We extend and apply SINDy to the case of learning reaction networks from non-equilibrium concentration data. Similar approaches make use of SINDy but do not resolve specific reactions [25], use weak formulations to avoid numerical temporal derivatives [28], or use compressive sensing and sparse Bayesian learning [27].

Our extension of the original SINDy method mostly involves estimating parameters which are coupled across the equations of the arising dynamical system. In the context of learning reaction networks this means that we look for specific reactions and their rate constants that might have lead to the observations instead of net flux across species. We demonstrate the algorithm on a gene regulatory network in three different scenarios of measurement: When there is no noise in the data we can find, given sufficient amounts of data, all relevant processes of the ground truth. If there is noise in the data we converge to the correct reaction network and rates with decreasing levels of noise. The final scenario generalizes the method to two measurements with different initial conditions, also converging to the correct model with decreasing levels of noise.

## II. REACTIVE SINDY: SPARSE LEARNING OF REACTION KINETICS

We are observing the concentrations of S chemical species in time *t*:

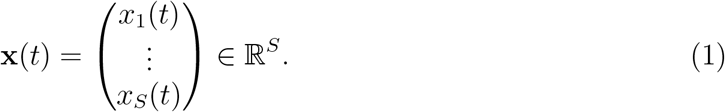

We assume that their dynamics are governed by classical reaction-rate equations subject to the law of mass action. A general expression for the change of concentration of reactant *s* as a result of order-0 reactions (creation or annihilation), order-1 reactions (transitions of other species into *s* or transitions of *s* into other species), order-2 reactions (production or consumption of *s* by the encounter of two species), etc, is given by:

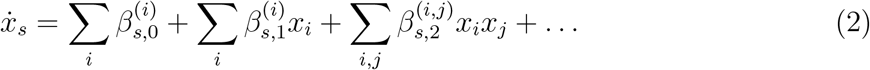

where the 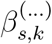-values are constants belonging to the reactions of order *k*. These rate constants however can incorporate several underlying reactions at once. For example, the two reactions

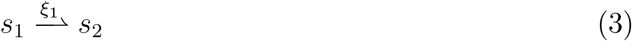

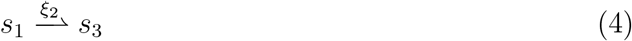

both contribute to 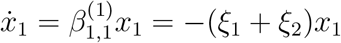. To disentangle (2) into single reactions, choose a library of *R* possible ansatz reactions that each represent a single reaction:

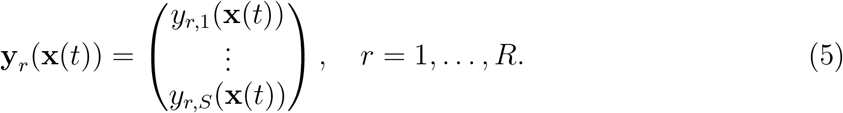

With this ansatz, the reaction dynamics (2) becomes a set of linear equations with unknown parameters *ξ*_*r*_ that represent the sought macroscopic rate constants:

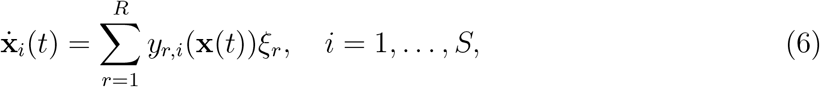

where *ξ*_*r*_ are the to-be estimated macroscopic rate constants. The two reactions in the previous example (3–4) would be modeled by the ansatz functions

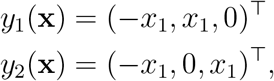

illustrating that the values of the coefficients *ξ*_1_ and *ξ*_2_ can be used to decide whether a single reaction is present and to what degree.

Now suppose we have measured the concentration vector (1) at *T* time points *t*_1_ < … < *t*_*T*_. We represent these data as a matrix

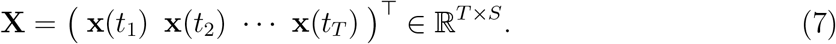

Given this matrix, a library Θ: ℝ^*T*×*S*^ → ℝ^*T*×*S*×*R*^, **X** ↦ (*θ*_1_(**X**) *θ*_2_(**X**) … *θ*_*R*_(**X**)) of *R* ansatz reactions can be proposed with corresponding reaction functions

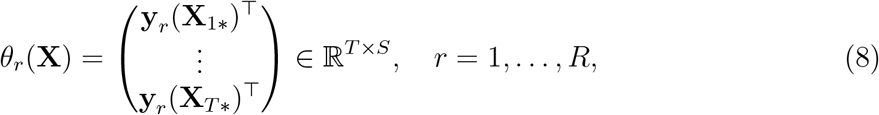

where **X**_*i*\*_ denotes the *i*-th row in **X**. Applying the concentration trajectory to the library yields Θ(**X**) ∊ ℝ^*T*×*S*×*R*^.

The goal is to find coefficients Ξ = (*ξ*_1_ *ξ*_2_ … *ξ*_*R*_)^T^, so that

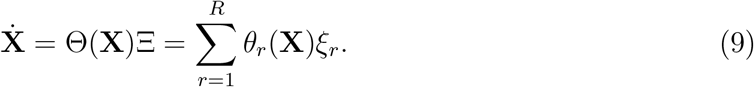

In particular, the system is linear in the coefficients Ξ, which makes regression tools such as elastic net regularization [45] applicable. To this end, one can consider the regularized minimization problem (reactive SINDy):

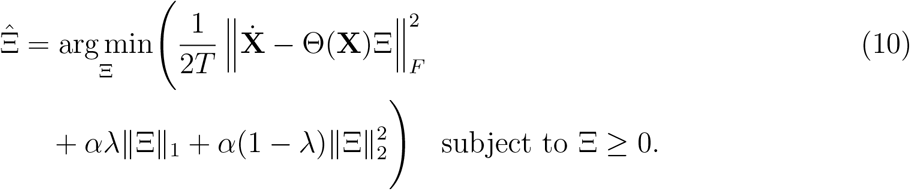

Here, ║ · ║_*F*_ denotes the Frobenius norm, λ ∊ [0,1] is a hyperparameter that interpolates linearly between LASSO [15, 38] and Ridge [16] methods, and α ≥ 0 is a hyperparameter that, depending on λ, can induce sparsity and give preference to smaller solutions in the *L*_1_ or *L*_2_ sense. For α = 0 the minimization problem reduces to standard least-squares (LSQ) with the constraint Ξ ≥ 0. Reactive SINDy (10) is therefore a generalization of the SINDy method to to the vector-valued ansatz functions.

Since only the concentration data **X** is available but not its temporal derivative, 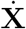 is approximated numerically by second order finite differences with the exception of boundary data. Once the pair 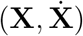 is obtained, the problem becomes invariant under temporal reordering. Hence, when presented with multiple trajectories the data matrices **X**_*i*_ and 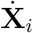 can simply be concatenated.

In order to solve (10) the numerical sequential least-squares minimizer SLSQP [21] is applied via the software package SciPy [19]. Code related to this paper can be found under https://github.com/readdy/readdy_learn.

## III. RESULTS

We demonstrate the method by estimating the reactions of a gene-regulatory network from time series of concentrations of the involved molecules. Let *S*: = {*A*,*B*,*C*} be a set of three species of proteins which are being translated each from their respective mRNA molecule. Each mRNA in turn has a corresponding DNA which it is transcribed from. The proteins and mRNA molecules decay over time whereas the DNA concentration remains constant. The network contains reactions of the following form [36]

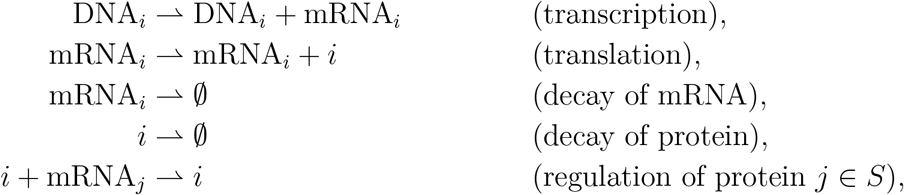

for each of the species *i* ∊ *S*. These reactions model a regulation of species *j* by virtue of the fact that the transcription product inhibits the transcription processes. In our example proteins of type A regulate the mRNA_B_ molecules, proteins of type B regulate the mRNA_C_ molecules and proteins of type C regulate the mRNA_A_ molecules (Fig. 1). Using this reaction model, time series of concentrations are generated using the rates given in Tab II under the initial condition described in Tab Ia, which were chosen so that all the reactions in the reaction model significantly contribute to the temporal evolution of the system’s concentrations. The generation samples the integrated equations equidistantly with a discrete time step of τ = 3 · 10^−3^ yielding 667 frames which amounts to a cumulative time of roughly *T* = 2.

**Figure 1.**
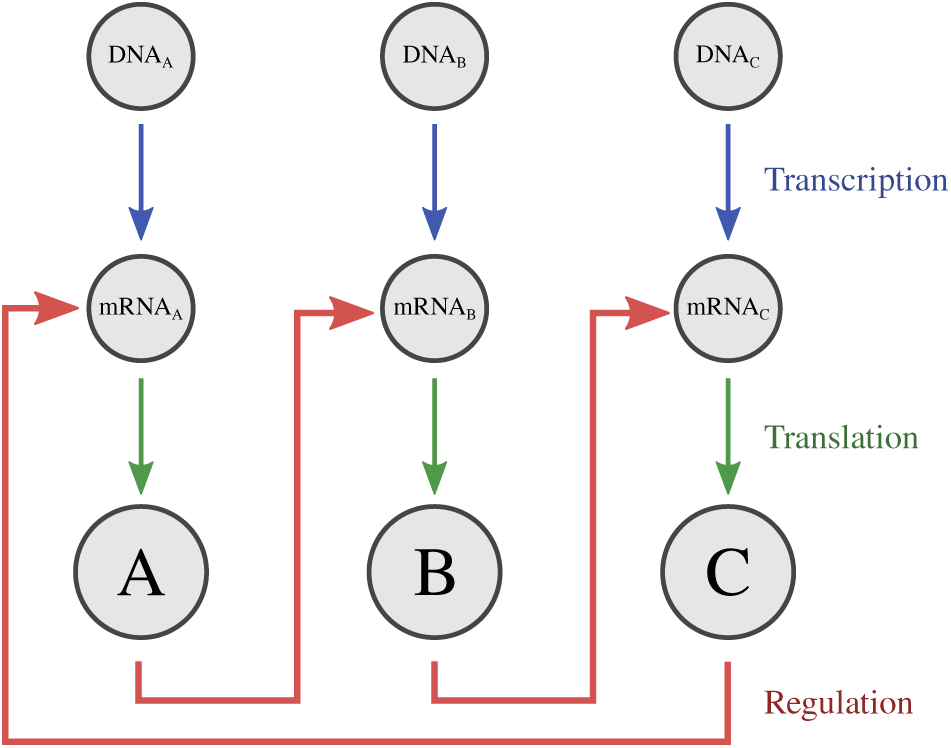
The regulation network example described in Sec. III. Each circle depicts a species, each arrow corresponds to one reaction. Blue arrows denote transcription from DNA to mRNA, green arrows denote translation from mRNA to protein, and red arrows denote the regulatory network.

**Table I.** Initial conditions **(a)** and **(b)** used to generate concentration time series. Reaction rates can be found in Tab. II.

|  | DNA<sub>A</sub> | mRNA<sub>A</sub> | A | DNA<sub>B</sub> | mRNA<sub>B</sub> | B | DNA<sub>C</sub> | mRNA<sub>C</sub> | C |
| --- | --- | --- | --- | --- | --- | --- | --- | --- | --- |
| (a) | 1 | 2 | 0 | 1 | 0 | 3 | 1 | 0 | 0 |
| (b) | 1 | 1.5 | 0 | 1 | 0 | 2 | 1 | 0 | 1 |

**Table II.**
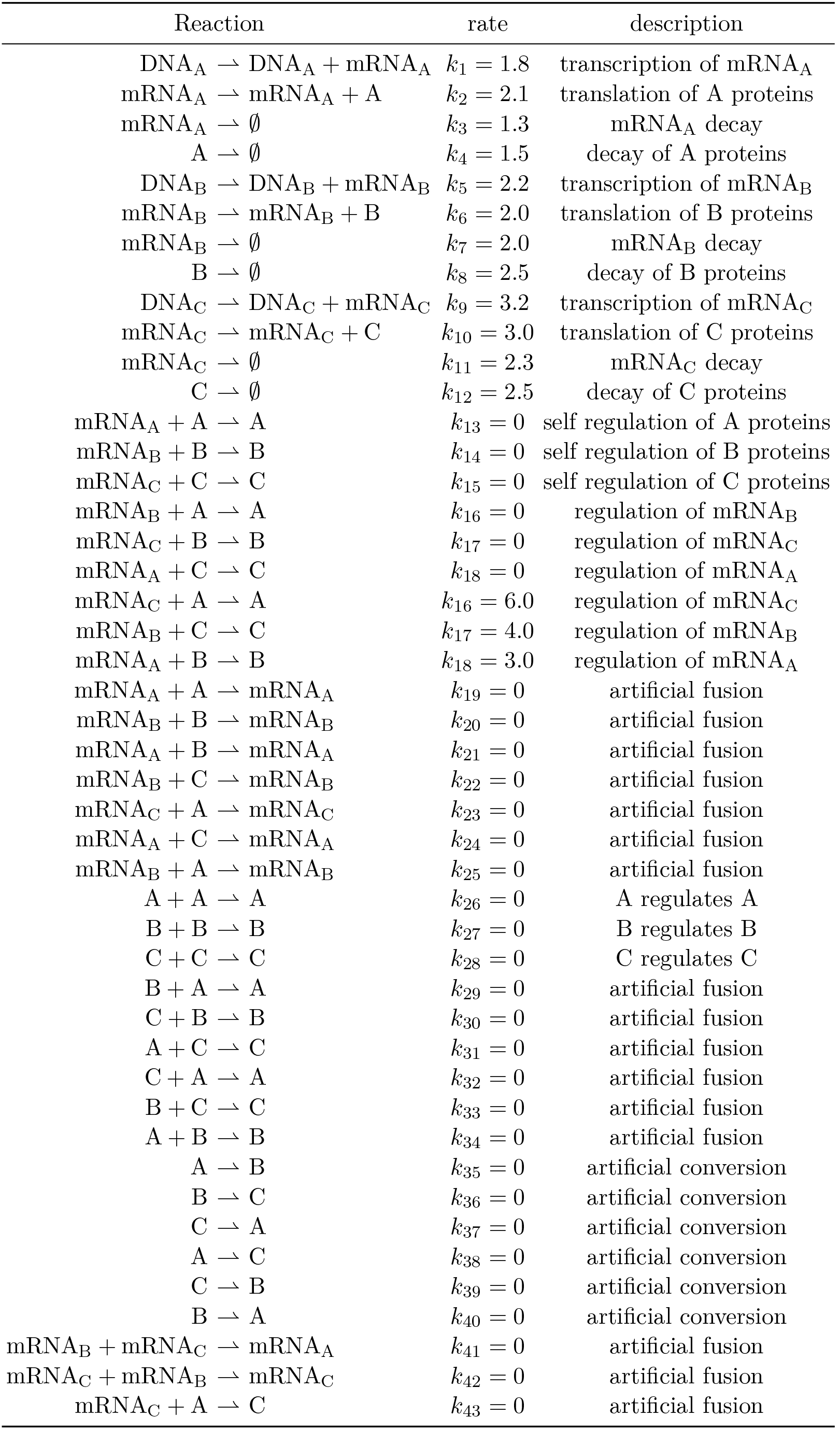
Full set of ansatz reactions Θ used in Sec. III. The given rate constants define the ground truth reaction model.

| Reaction | rate | description |
| --- | --- | --- |
| $\text{DNA}_A \rightarrow \text{DNA}_A + \text{mRNA}_A$ | $k_1 = 1.8$ | transcription of $\text{mRNA}_A$ |
| $\text{mRNA}_A \rightarrow \text{mRNA}_A + A$ | $k_2 = 2.1$ | translation of A proteins |
| $\text{mRNA}_A \rightarrow \emptyset$ | $k_3 = 1.3$ | $\text{mRNA}_A$ decay |
| $A \rightarrow \emptyset$ | $k_4 = 1.5$ | decay of A proteins |
| $\text{DNA}_B \rightarrow \text{DNA}_B + \text{mRNA}_B$ | $k_5 = 2.2$ | transcription of $\text{mRNA}_B$ |
| $\text{mRNA}_B \rightarrow \text{mRNA}_B + B$ | $k_6 = 2.0$ | translation of B proteins |
| $\text{mRNA}_B \rightarrow \emptyset$ | $k_7 = 2.0$ | $\text{mRNA}_B$ decay |
| $B \rightarrow \emptyset$ | $k_8 = 2.5$ | decay of B proteins |
| $\text{DNA}_C \rightarrow \text{DNA}_C + \text{mRNA}_C$ | $k_9 = 3.2$ | transcription of $\text{mRNA}_C$ |
| $\text{mRNA}_C \rightarrow \text{mRNA}_C + C$ | $k_{10} = 3.0$ | translation of C proteins |
| $\text{mRNA}_C \rightarrow \emptyset$ | $k_{11} = 2.3$ | $\text{mRNA}_C$ decay |
| $C \rightarrow \emptyset$ | $k_{12} = 2.5$ | decay of C proteins |
| $\text{mRNA}_A + A \rightarrow A$ | $k_{13} = 0$ | self regulation of A proteins |
| $\text{mRNA}_B + B \rightarrow B$ | $k_{14} = 0$ | self regulation of B proteins |
| $\text{mRNA}_C + C \rightarrow C$ | $k_{15} = 0$ | self regulation of C proteins |
| $\text{mRNA}_B + A \rightarrow A$ | $k_{16} = 0$ | regulation of $\text{mRNA}_B$ |
| $\text{mRNA}_C + B \rightarrow B$ | $k_{17} = 0$ | regulation of $\text{mRNA}_C$ |
| $\text{mRNA}_A + C \rightarrow C$ | $k_{18} = 0$ | regulation of $\text{mRNA}_A$ |
| $\text{mRNA}_C + A \rightarrow A$ | $k_{16} = 6.0$ | regulation of $\text{mRNA}_C$ |
| $\text{mRNA}_B + C \rightarrow C$ | $k_{17} = 4.0$ | regulation of $\text{mRNA}_B$ |
| $\text{mRNA}_A + B \rightarrow B$ | $k_{18} = 3.0$ | regulation of $\text{mRNA}_A$ |
| $\text{mRNA}_A + A \rightarrow \text{mRNA}_A$ | $k_{19} = 0$ | artificial fusion |
| $\text{mRNA}_B + B \rightarrow \text{mRNA}_B$ | $k_{20} = 0$ | artificial fusion |
| $\text{mRNA}_A + B \rightarrow \text{mRNA}_A$ | $k_{21} = 0$ | artificial fusion |
| $\text{mRNA}_B + C \rightarrow \text{mRNA}_B$ | $k_{22} = 0$ | artificial fusion |
| $\text{mRNA}_C + A \rightarrow \text{mRNA}_C$ | $k_{23} = 0$ | artificial fusion |
| $\text{mRNA}_A + C \rightarrow \text{mRNA}_A$ | $k_{24} = 0$ | artificial fusion |
| $\text{mRNA}_B + A \rightarrow \text{mRNA}_B$ | $k_{25} = 0$ | artificial fusion |
| $A + A \rightarrow A$ | $k_{26} = 0$ | A regulates A |
| $B + B \rightarrow B$ | $k_{27} = 0$ | B regulates B |
| $C + C \rightarrow C$ | $k_{28} = 0$ | C regulates C |
| $B + A \rightarrow A$ | $k_{29} = 0$ | artificial fusion |
| $C + B \rightarrow B$ | $k_{30} = 0$ | artificial fusion |
| $A + C \rightarrow C$ | $k_{31} = 0$ | artificial fusion |
| $C + A \rightarrow A$ | $k_{32} = 0$ | artificial fusion |
| $B + C \rightarrow C$ | $k_{33} = 0$ | artificial fusion |
| $A + B \rightarrow B$ | $k_{34} = 0$ | artificial fusion |
| $A \rightarrow B$ | $k_{35} = 0$ | artificial conversion |
| $B \rightarrow C$ | $k_{36} = 0$ | artificial conversion |
| $C \rightarrow A$ | $k_{37} = 0$ | artificial conversion |
| $A \rightarrow C$ | $k_{38} = 0$ | artificial conversion |
| $C \rightarrow B$ | $k_{39} = 0$ | artificial conversion |
| $B \rightarrow A$ | $k_{40} = 0$ | artificial conversion |
| $\text{mRNA}_B + \text{mRNA}_C \rightarrow \text{mRNA}_A$ | $k_{41} = 0$ | artificial fusion |
| $\text{mRNA}_C + \text{mRNA}_B \rightarrow \text{mRNA}_C$ | $k_{42} = 0$ | artificial fusion |
| $\text{mRNA}_C + A \rightarrow C$ | $k_{43} = 0$ | artificial fusion |

The proposed estimation method is applied to analyze these time series of concentrations in order to recover the underlying reaction network from data. To this end we use the library of ansatz functions given in Tab. II, which contains a large number of possible reactions, only few of which are actually part of the model.

### A. Learning the reaction network in the low-noise limit

We first demonstrate that the true reaction network can be reconstructed when using a finite amount of observation data without additional measurement noise, i.e., the observations are reflecting the true molecule concentrations at any given time point. The minimization problem (10) is solved using the concentration time series shown in Fig. 1b.

We first set the hyperparameter α = 0 in the minimization problem (10), which results in constrained least-squares regression without any of the regularization terms. In this case we estimate a reaction network that can reproduce the observations almost exactly (Fig. 2). However, the result is mechanistically wrong as the sparsity pattern does not match the reaction network used to generate the data. On the one hand many spurious reactions are estimated that were not in the true reaction scheme and would lead to wrong conclusions about the mechanism, such as A + A ⇀ A and A + C ⇀ C. More dramatically, the reaction responsible for the decay of A particles is completely ignored (Fig. 3).

**Figure 2.**
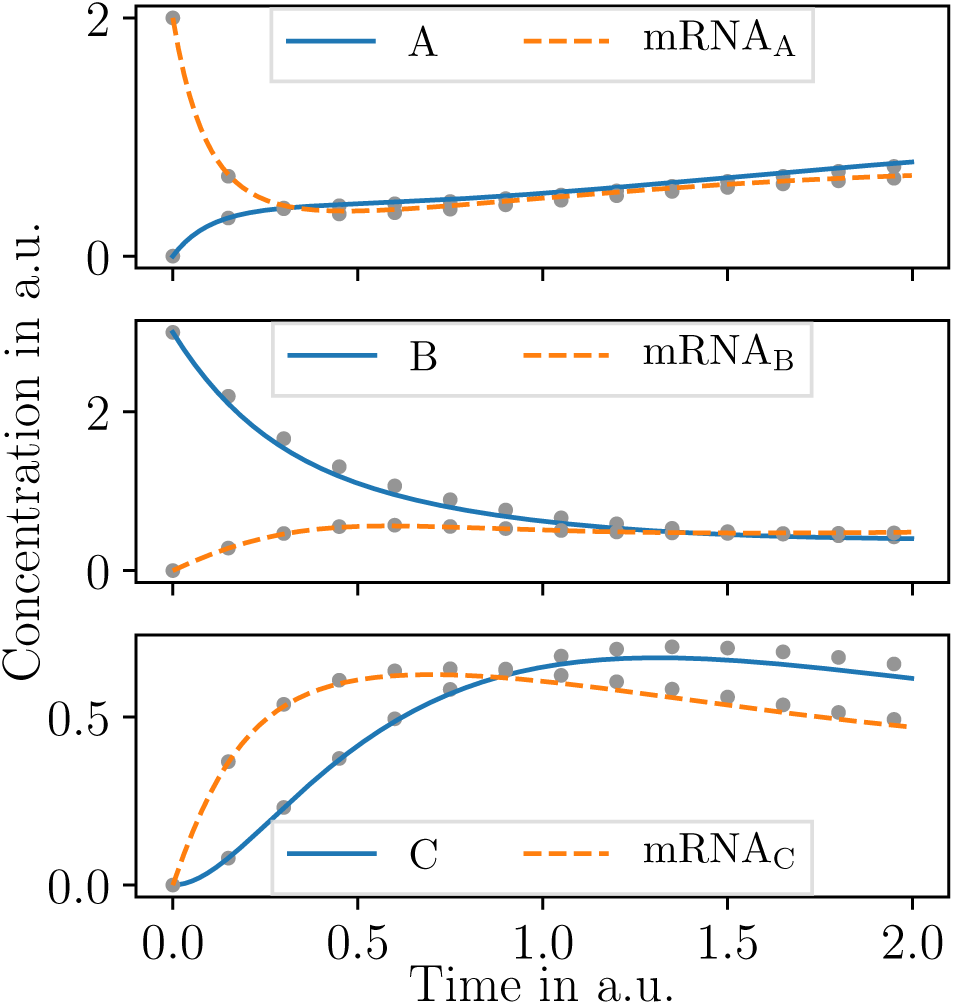
Concentration time series generated from integrating the reaction network shown in Fig. 1a. The initial condition prescribes positive concentration values only for B protein and mRNAA species (Tab. Ia). This initial condition is used in the subsequent sections for further analysis. Gray dots depict concentration time series yielded from the LSQ rates estimated in Sec III A.

**Figure 3.**
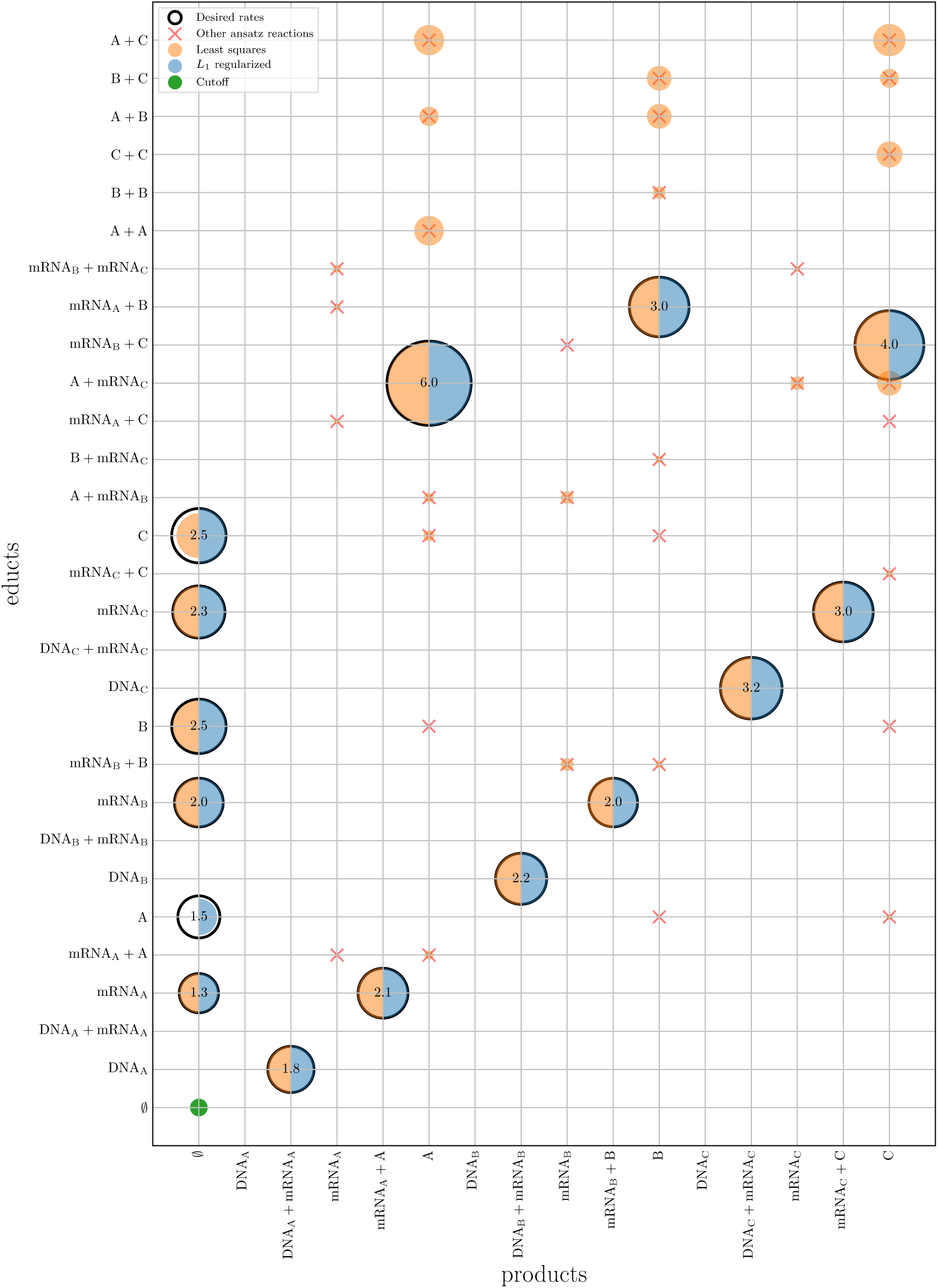
Estimated reaction rates in the system described in Sec. III A. The y and x axes contain reaction educts and products, respectively. A circle at position (*i*, *j*) represents a reaction *i* ⇀ *j* whose rate has a linear relation with the area of the circle. The black outlines denote the reactions with which the system was generated and contain the respective rate value. Red crosses denote reactions that were used as additional ansatz reactions. Blue circles are estimated by LSQ and orange circles depict rates which were obtained by solving the minimization problem (10). The latter rates are subject to a cutoff κ = 0.22 corresponding to the green circle’s area under which a sparse solution with the correct processes can be recovered. If a certain rate was estimated in both cases, two wedges instead of one circle are displayed.

Next, we sought sparse solutions by using α > 0 and additionally eliminating reactions with rate constants smaller than a cutoff value κ. For a suitable choice of hyperparameters α ≈ 1.91 · 10^−7^, λ =1, and κ = 0.22, a sparse solution is obtained that finds the correct reaction scheme and also recovers the decay reaction (Fig. 3).

The value of the cutoff was determined by comparing the magnitude of estimated rates and finding a gap, see Fig. 6. The hyperparameter pair (α, λ) was obtained by a grid search and evaluating the difference 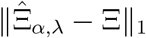, where 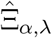 is the estimated model under a particular hyperparameter choice and Ξ is the ground truth. If the ground truth is unknown, a hyperparameter pair can be estimated by utilizing cross-validation as in the following sections.

### B. Learning the reaction network from data with stochastic noise

In contrast to Sec. III A, we now employ data that includes measurement noise. Such noise can originate from uncertainties in the experimental setup or from shot noise in single-or few-molecule measurements. In gene regulatory networks such noise is commonly observed when only few copy numbers of mRNA are present [4, 9, 14]. In order to simulate noise from few copies of molecules, the system of Sec. III with initial conditions as given in Tab. Ia is integrated using the Gillespie stochastic simulation algorithm (SSA) [12, 13]. In the limit of many particles and realizations, the Gillespie SSA converges to the integrated reaction-rate equations subject to the law of mass action. As our model is based on exactly these dynamics, the initial condition’s concentrations are interpreted in terms of hundreds of particles. Each realization is then transformed back to a time series of concentrations. We define the noise level as the mean-squared deviation of the concentration time series from the integrated reaction-rate equations. Data with different noise levels are prepared by averaging multiple realizations of the time series obtained by the Gillespie SSA.

It can be observed that decreasing levels of noise lead to fewer spurious reactions when applying reactive SINDy (10), see Fig. 4a. Also the estimation error 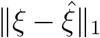 with respect to the ground truth *ξ* decreases with decreasing levels of noise (Fig. 4b). In both cases, the regularized method with a suitable hyperparameter pair (α, λ) performs better than LSQ.

**Figure 4.**
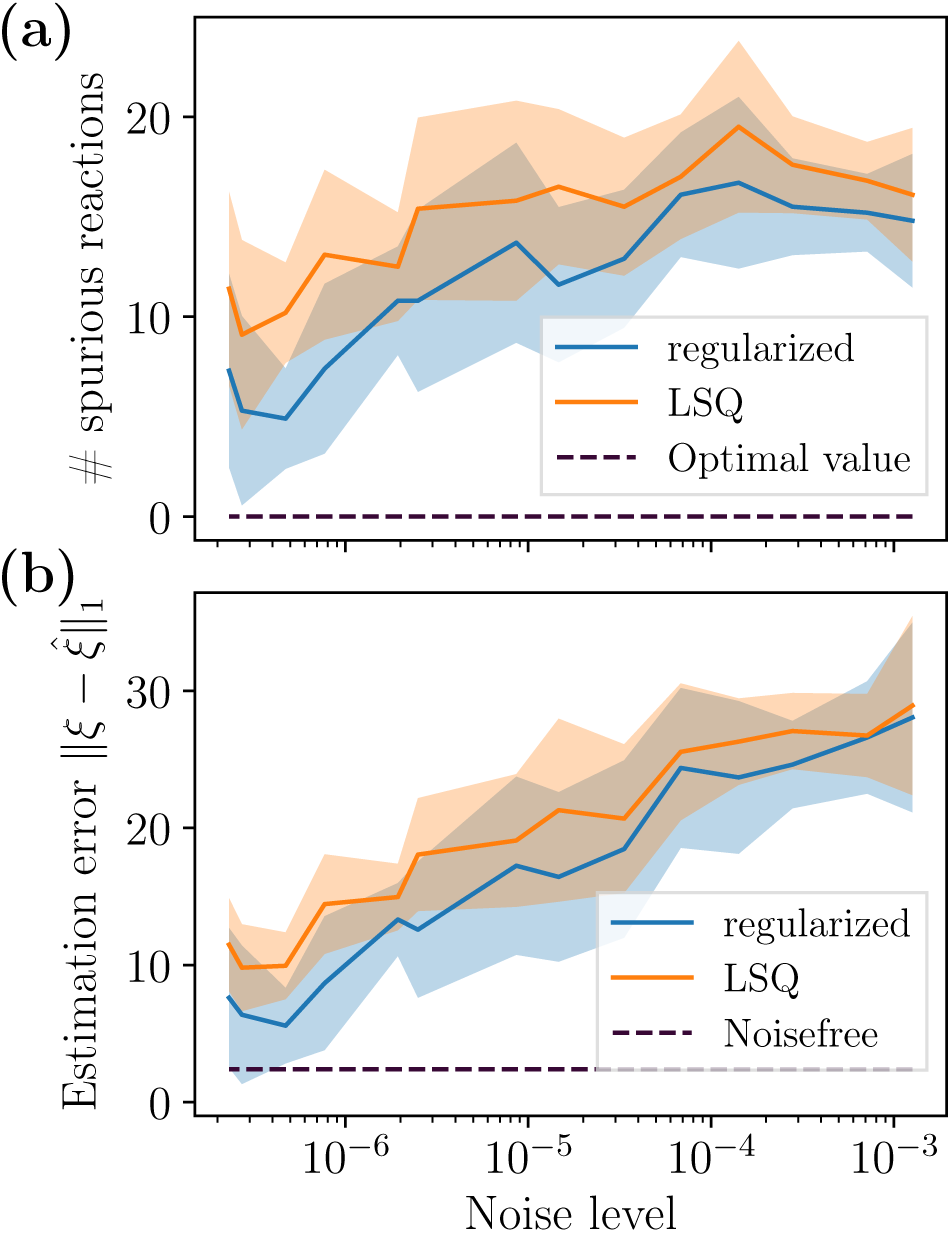
Convergence of the estimation error when estimating the system described in Sec. IIIA with varying levels of noise by application of reactive SINDy (10) with and without regularization in blue and orange, respectively. The procedure was independently repeated 10 times with different realizations giving rise to the mean and standard deviation depicted by solid lines and shaded areas, respectively. **(a)**: The number of detected spurious reactions up to the cutoff value introduced in Sec. IIIA over different levels of noise. **(b)**: The estimation error given by the mean absolute error between the generating reaction rates *ξ* and the estimated reaction rates 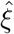 over different levels of noise.

The hyperparameters (α, λ) are obtained by shuffling the data and performing a 10-fold cross validation.

### C. Learning the reaction network from multiple initial conditions

Preparing the experiment that generates the data in different initial conditions can help identifying the true reaction mechanisms as a more diverse dataset makes it easier to confirm or exclude the participation of specific reactions. This section extends the analysis of Sec. IIIB to two initial conditions, where the first initial condition is identical to the one used previously and the second initial condition is given in Tab. Ib.

The corresponding time series are depicted in Fig. 5a. The gray graph corresponds to a sample trajectory generated by the Gillespie SSA. For both initial conditions the same time step of *τ* = 3 · 10^−3^ has been applied, amounting to 2 · 667 = 1334 frames. Once the data matrices

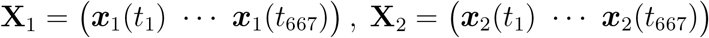

and the corresponding derivatives 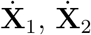 have been obtained, the frames are concatenated so that

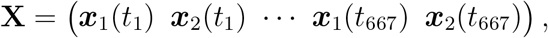

analogously for 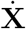.

**Figure 5.**
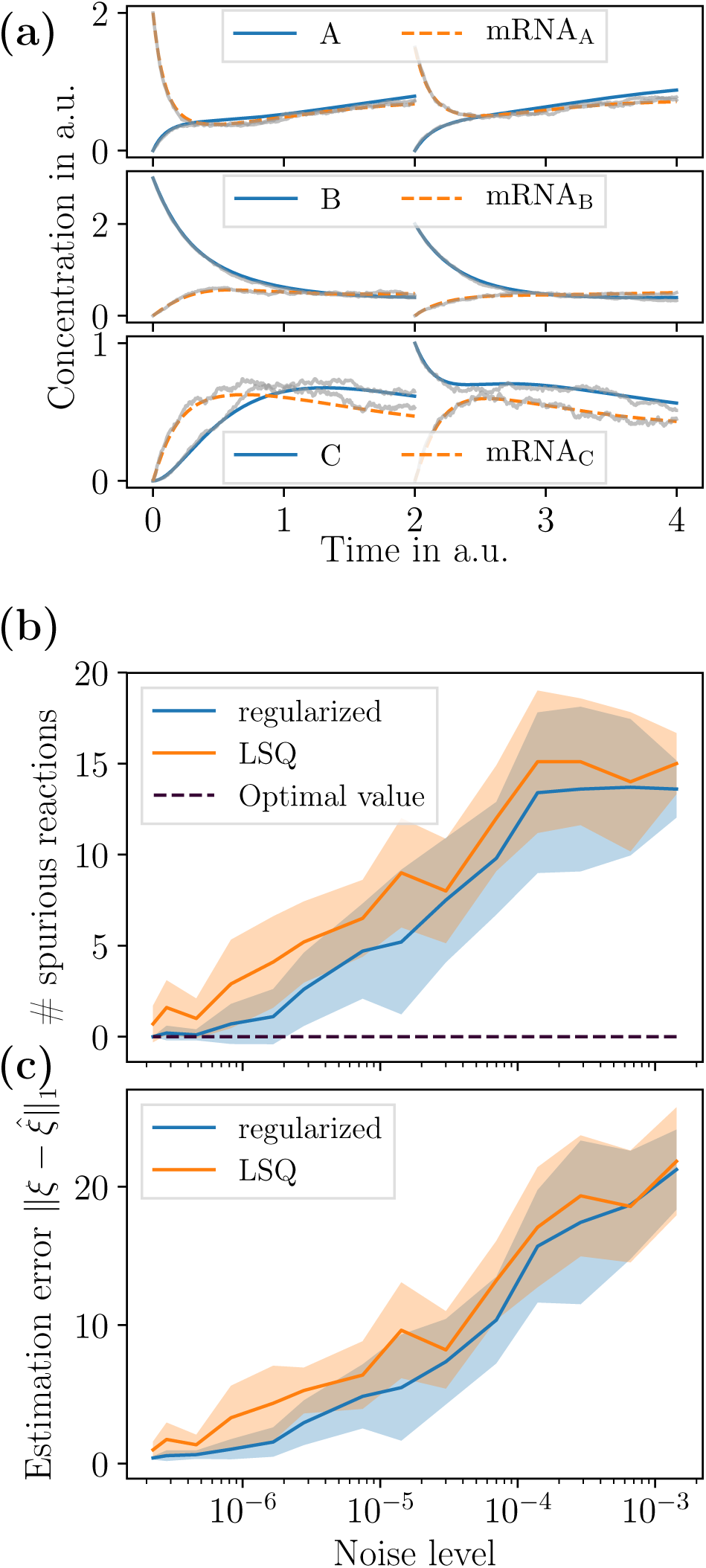
Convergence of estimation error of reaction schemes from noisy gene-regulation data starting from two different initial conditions under decreasing levels of noise. The minimization problem (10) was solved for α = 0 (LSQ) and with regularization. This was repeated 10 times on different sets of observation data generated by Gillespie SSA, giving rise to mean and standard deviation (solid lines and shaded areas, respectively). **(a)**: Concentration time series corresponding to the initial conditions, generated by integrating the reaction-rate equations. The first initial condition is identical to the one used in Sec. IIIA and Sec. IIIB. The second initial condition (Tab. Ib) prescribes positive initial concentrations for mRNA_A_, B, and C species. The gray graphs are sample realizations of integration using the Gillespie SSA. **(b)**,**(c)**: Analogously to Fig. 4 with the difference that 20-fold cross validation was used for hyperparameter estimation.

Similarly to Sec. IIIB, decreasing levels of noise lead to fewer spurious reactions (Fig. 5b) and a smaller *L*_1_ distance to the ground truth (Fig. 5c). Again applying the optimization problem with a suitable set of parameters (α, λ, κ) performs better than LSQ. Compared to the previous section the convergence is better due to twice as much available data. At noise levels of smaller than roughly 10^−6^ the model can reliably be recovered when using the regularized method.

The hyperparameters (α, λ) are obtained by shuffling the data and performing a 20-fold cross validation.

## IV. CONCLUSION

In this work we have extended the SINDy method to reactive SINDy, not only parsimoniously detecting potentially nonlinear terms in a dynamical system from noisy data, but also yielding, in this case, a sparse set of rates with respect to generating reactions(8). Mathematically this has been achieved by permitting vector-valued basis functions and obtaining a tensor linear regression problem. We have applied this method on data generated from a gene regulation network and could successfully recover the underlying model.

The studies of Sec. IIIB and Sec. IIIC have shown that the applied regularization terms can mitigate noise up to a certain degree compared to the unregularized method, so that identification of the reaction network is more robust and closer to the ground truth.

Potentially, this method could be used to identify reaction networks from time series measurements even if the initial conditions are not always exactly identical, as was demonstrated in Sec. IIIC. One apparent limitation is that the method can only be applied if the data stems from the equilibration phase, as the concentration-based approach has derivatives equal zero in the equilibrium, which precludes the reaction dynamics to be recovered. In future work, we will consider the identification of reaction schemes from instantaneous fluctuations of particle numbers in equilibrium.

## ACKNOWLEDGMENTS

The authors are grateful to the Center for Theoretical Biological Physics (CTBP, supported by NSF PHY-1427654) at Rice University for hosting their sabbatical visit, during which part of this work was performed.

We gratefully acknowledge funding from Deutsche Forschungsgemeinschaft (SFB 958 / Project A04, TRR 186 / Project A12, SFB 1114 / Project C03), Einstein Foundation Berlin (ECMath Project CH17) and European Research Council (ERC CoG 772230 “ScaleCell”). We are grateful for inspiring discussions with Simon Olsson, Mohsen Sadeghi, Felix Höfling and Christof Schütte.

## V. APPENDIX

**Figure 6.**
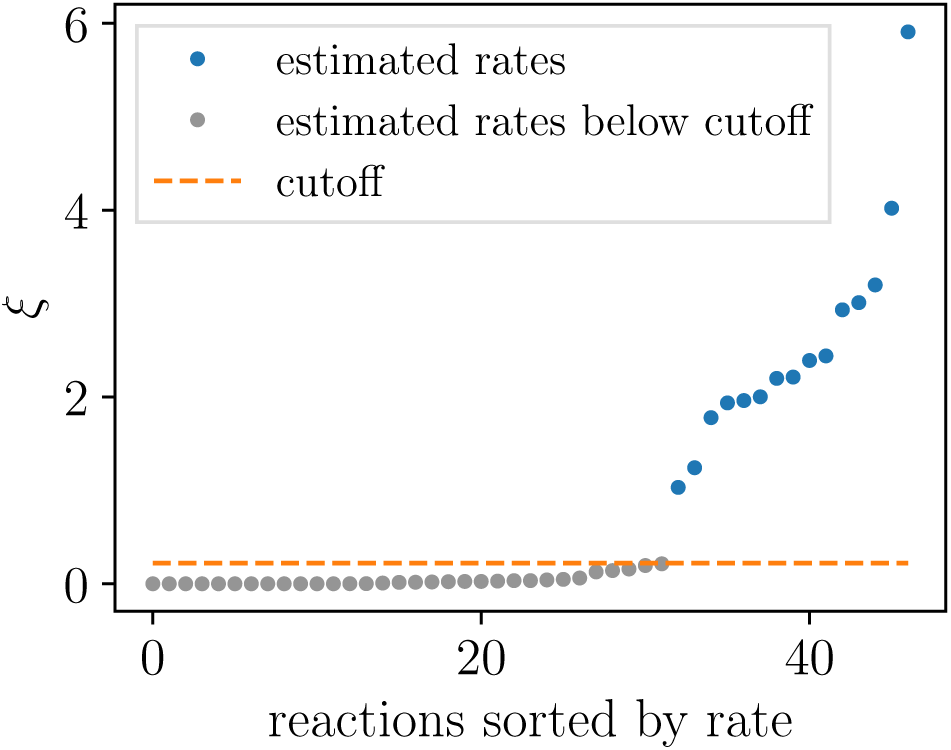
Reaction rates sorted by their magnitude to determine the cutoff *κ* = 0.22 of Sec. III A. The rates were estimated using the regularized minimization problem.

## References

[1] R. Abramovitch, E. Tavor, J. Jacob-Hirsch, E. Zeira, N. Amariglio, O. Pappo, G. Rechavi, E. Galun, and A. Honigman. A Pivotal Role of Cyclic AMP-Responsive Element Binding Protein in Tumor Progression. Cancer Research, 64(4):1338–1346, feb 2004. I

[2] S. S. Andrews. Smoldyn: Particle-based simulation with rule-based modeling, improved molecular interaction and a library interface. Bioinformatics, 33(5):710–717, 2017. I

[3] S. S. Andrews. Particle-Based Stochastic Simulators, pages 1–5. Springer New York, New York, NY, 2018. I

[4] O. G. Berg. A model for the statistical fluctuations of proteins numbers in a microbial population. J. Theor. Biol., 71(4):587–603, 1978. III B

[5] S. L. Brunton, J. L. Proctor, and J. N. Kutz. Discovering governing equations from data by sparse identification of nonlinear dynamical systems. PNAS, 1(609):1–26, 2015. I

[6] S. L. Brunton, J. L. Proctor, and J. N. Kutz. Sparse identification of nonlinear dynamics with control (sindyc). arXiv preprint arXiv:1605.06682, 2016. I

[7] P. Cong, R. D. Doolen, Q. Fan, D. M. Giaquinta, S. Guan, E. W. McFarland, D. M. Poojary, K. Self, H. W. Turner, and W. H. Weinberg. High-throughput synthesis and screening of combinatorial heterogeneous catalyst libraries. Angewandte Chemie International Edition, 38(4):483–488, 1999. I

[8] A. Donev, C.-Y. Yang, and C. Kim. Efficient reactive brownian dynamics. The Journal of chemical physics, 148(3):034103, 2018. I

[9] M. B. Elowitz. Stochastic Gene Expression in a Single Cell. Science, 297(5584):1183–1186, aug 2002. III B

[10] C. Fröhner and F. Noé. Reversible interacting-particle reaction dynamics. The Journal of Physical Chemistry B, 2018. I

[11] S. Gama-Castro, H. Salgado, A. Santos-Zavaleta, D. Ledezma-Tejeida, L. Muñiz-Rascado, J. S. García-Sotelo, K. Alquicira-Hernández, I. Martínez-Flores, L. Pannier, J. A. Castro-Mondragón, A. Medina-Rivera, H. Solano-Lira, C. Bonavides-Martínez, E. Pérez-Rueda, S. Alquicira-Hernández, L. Porrón-Sotelo, A. López-Fuentes, A. Hernández-Koutoucheva, V. D. Moral-Chávez, F. Rinaldi, and J. Collado-Vides. RegulonDB version 9.0: high-level integration of gene regulation, coexpression, motif clustering and beyond. Nucleic Acids Research, 44(D1):D133–D143, jan 2016. I

[12] D. T. Gillespie. A general method for numerically simulating the stochastic time evolution of coupled chemical reactions. Journal of computational physics, 22(4):403–434, 1976. IIIB

[13] D. T. Gillespie. Exact stochastic simulation of coupled chemical reactions. The journal of physical chemistry, 81(25):2340–2361, 1977. I, IIIB

[14] I. Golding, J. Paulsson, S. M. Zawilski, and E. C. Cox. Real-time kinetics of gene activity in individual bacteria. Cell, 123(6):1025–1036, 2005. IIIB

[15] T. Hastie, R. Tibshirani, and J. Friedman. The Elements of Statistical Learning: Data Mining, Inference, and Prediction. Springer New York, New York, NY, 2009. II

[16] A. E. Hoerl and R. W. Kennard. Ridge regression: Biased estimation for nonorthogonal problems. Technometrics, 12:55–67, 1970. II

[17] M. Hoffmann, C. Fröhner, and F. Noé. Readdy 2: Fast and flexible software framework for interacting-particle reaction dynamics. bioRxiv, page 374942, 2018. I

[18] S. A. Isaacson. The reaction-diffusion master equation as an asymptotic approximation of diffusion to a small target. SIAM Journal on Applied Mathematics, 70(1):77–111, 2009. I

[19] E. Jones, T. Oliphant, P. Peterson, et al. SciPy: Open source scientific tools for Python, 2001-. [Online; accessed October 27, 2017]. II

[20] L. Kiwi-Minsker and A. Renken. Microstructured reactors for catalytic reactions. Catalysis today, 110(1-2):2–14, 2005. I

[21] D. Kraft. A software package for sequential quadratic programming. Technical Report DFVLR-FB 88-28, Institut für Dynamik der Flugsysteme, Oberpfaffenhofen, 1988. II

[22] T. Kuhlman, Z. Zhang, M. H. Saier, and T. Hwa. Combinatorial transcriptional control of the lactose operon of Escherichia coli. Proceedings of the National Academy of Sciences, 104(14):6043–6048, apr 2007. I

[23] T. I. Lee. Transcriptional Regulatory Networks in Saccharomyces cerevisiae. Science, 298(5594):799–804, oct 2002. I

[24] M. Lewis, G. Chang, N. C. Horton, M. A. Kercher, H. C. Pace, M. A. Schumacher, R. G. Brennan, and P. Lu. Crystal Structure of the Lactose Operon Repressor and Its Complexes with DNA and Inducer. Science, 271(5253):1247–1254, mar 1996. I

[25] N. M. Mangan, S. L. Brunton, J. L. Proctor, and J. N. Kutz. Inferring biological networks by sparse identification of nonlinear dynamics. IEEE Transactions on Molecular, Biological and Multi-Scale Communications, 2(1):52–63, 2016. I

[26] A. A. Margolin, I. Nemenman, K. Basso, C. Wiggins, G. Stolovitzky, R. Favera, and A. Califano. ARACNE: An Algorithm for the Reconstruction of Gene Regulatory Networks in a Mammalian Cellular Context. BMC Bioinformatics, 7(Suppl 1):S7, 2006. I

[27] W. Pan, Y. Yuan, J. Goncalves, and G.-b. Stan. Reconstruction of arbitrary biochemical reaction networks: A compressive sensing approach. In 2012 IEEE 51st IEEE Conference on Decision and Control (CDC), pages 2334–2339. IEEE, dec 2012. I

[28] Y. Pantazis and I. Tsamardinos. A unified approach for sparse dynamical system inference from temporal measurements. arXiv preprint arXiv:1710.00718, 2017. I

[29] M. Quade, M. Abel, J. Nathan Kutz, and S. L. Brunton. Sparse identification of nonlinear dynamics for rapid model recovery. Chaos: An Interdisciplinary Journal of Nonlinear Science, 28(6):063116, 2018. I

[30] R. Roa, W. K. Kim, M. Kandu, J. Dzubiella, and S. Angioletti-Uberti. Catalyzed Bimolecular Reactions in Responsive Nanoreactors. ACS Catalysis, 7(9):56045611, sep 2017. I

[31] S. H. Rudy, S. L. Brunton, J. L. Proctor, and J. N. Kutz. Data-driven discovery of partial differential equations. Science Advances, 3(4):e1602614, 2017. I

[32] J. Schöneberg, M. Heck, K. P. Hofmann, and F. Noé. Explicit spatiotemporal simulation of receptor-g protein coupling in rod cell disk membranes. Biophysical journal, 107(5):1042–1053, 2014. I

[33] J. Schöneberg and F. Noé. Readdy-a software for particle-based reaction-diffusion dynamics in crowded cellular environments. PLoS One, 8(9):e74261, 2013. I

[34] J. Schöneberg, A. Ullrich, and F. Noé. Simulation tools for particle-based reaction-diffusion dynamics in continuous space. BMC biophysics, 7(1):11, 2014. I

[35] S. S. Shen-Orr, R. Milo, S. Mangan, and U. Alon. Network motifs in the transcriptional regulation network of Escherichia coli. Nature Genetics, 31(1):64–68, may 2002. I

[36] M. Thattai and A. van Oudenaarden. Intrinsic noise in gene regulatory networks. Proceedings of the National Academy of Sciences, 98(15):8614–8619, 2001. III

[37] T. Tian, S. Xu, J. Gao, and K. Burrage. Simulated maximum likelihood method for estimating kinetic rates in gene expression. Bioinformatics, 23(1):84–91, 2007. I

[38] R. Tibshirani. Regression Selection and Shrinkage via the Lasso, 1996. II

[39] J. S. van Zon and P. R. Ten Wolde. Greens-function reaction dynamics: a particle-based approach for simulating biochemical networks in time and space. The Journal of chemical physics, 123(23):234910, 2005. I

[40] J. S. van Zon and P. R. Ten Wolde. Simulating biochemical networks at the particle level and in time and space: Greens function reaction dynamics. Physical review letters, 94(12):128103, 2005. I

[41] S. Winkelmann and C. Schütte. The spatiotemporal master equation: Approximation of reaction-diffusion dynamics via markov state modeling. The Journal of Chemical Physics, 145(21):214107, 2016. I

[42] S. Winkelmann and C. Schütte. Hybrid models for chemical reaction networks: Multiscale theory and application to gene regulatory systems. The Journal of chemical physics, 147(11):114115, 2017. I

[43] N. Yildirim and M. C. Mackey. Feedback Regulation in the Lactose Operon: A Mathematical Modeling Study and Comparison with Experimental Data. Biophysical Journal, 84(5):2841–2851, may 2003. I

[44] L. Zhang and H. Schaeffer. On the convergence of the sindy algorithm. arXiv preprint arXiv:1805.06445, 2018. I

[45] H. Zou and T. Hastie. Regularization and variable selection via the elastic-net. Journal of the Royal Statistical Society, 67(2):301–320, 2005. II

